# TDP-43 oligomers detected as initial intermediate species during aggregate formation

**DOI:** 10.1101/499343

**Authors:** Rachel L. French, Ashley N. Reeb, Himani Aligireddy, Niraja Kedia, Dhruva D. Dhavale, Zachary R. Grese, Paul T. Kotzbauer, Jan Bieschke, Yuna M. Ayala

## Abstract

Aggregates of the RNA binding protein TDP-43 are a hallmark of amyotrophic lateral sclerosis (ALS) and frontotemporal dementia (FTD), which are neurodegenerative disorders with overlapping clinical, genetic and pathological features. Mutations in the TDP-43 gene are causative of ALS, supporting its central role in pathogenesis. The process of TDP-43 aggregation remains poorly understood and whether this includes formation of intermediate complexes is unknown. We characterized aggregates derived from purified TDP-43 as a function of time and analyzed them under semi-denaturing conditions. Our assays identified oligomeric complexes at the initial time points prior to the formation of large aggregates, suggesting that ordered oligomerization is an intermediate step of TDP-43 aggregation. In addition, we analyzed liquid-liquid phase separation of TDP-43 and detected similar oligomeric assembly upon the maturation of liquid droplets into solid-like fibrils. These results strongly suggest that the oligomers form during the early steps of TDP-43 misfolding. Importantly, ALS-linked mutations A315T and M337V significantly accelerate aggregation, rapidly decreasing the monomeric population and shortening the oligomeric phase. We also show that the aggregates generated from purified protein seed intracellular aggregation, which is detected by established markers of TDP-43 pathology. Remarkably, cytoplasmic aggregate propagation is detected earlier with A315T and M337V and is 50% more widespread than with wild-type aggregates. Our findings provide evidence for a controlled process of TDP-43 self-assembly into intermediate structures that provide a scaffold for aggregation. This process is altered by ALS-linked mutations, underscoring the role of perturbations in TDP-43 homeostasis in protein aggregation and ALS-FTD pathogenesis.

## INTRODUCTION

TDP-43 pathology is a hallmark of amyotrophic lateral sclerosis (ALS) and frontotemporal dementia (FTD). TDP-43 aggregates accumulate in approximately 98% of ALS and 50% of FTD cases, also defined as ubiquitin-positive frontotemporal lobar degeneration (FTLD-U or FTLD-TDP) (1,2). In addition, TDP-43 pathology is found in approximately 50% of Alzheimer’s disease (3,4). The direct role of TDP-43 in disease is underscored by greater than 40 ALS-associated dominant missense mutations in the TDP-43 gene (*TARDBP*) (5). These cause approximately 3-5% and 1% of familial and sporadic ALS, respectively. The clinico-pathological characteristics of ALS and FTD associated with mutant and wild-type *TARDBP* are largely indistinguishable, and the mechanisms affected by the mutations linked to pathogenesis have not been clearly established. Whether disease results from gain of toxic properties through aggregation, from sequestration of functional TDP-43 into aggregates (1), or from a combination of both, it is increasingly evident that loss of TDP-43 homeostasis and aggregation play a critical role in pathogenesis.

TDP-43 is a highly conserved RNA binding protein and, as other heterogeneous nuclear ribonucleoproteins (hnRNPs), is composed of modular domains that mediate single stranded RNA/DNA binding and protein interactions (6–8). Of the two canonical RNA recognition motifs (RRMs), RRM1 contributes to the high affinity for RNA/DNA and GU-rich RNA specificity (6,7). RRM2 is also highly evolutionarily conserved, however, its function remains unclear. An additional folded domain is at the amino terminus, which mediates self-assembly as an isolated domain and presumably of the full-length protein (9–11). The carboxy-terminal domain (CTD) is intrinsically disordered and is a typical low sequence complexity domain (LCD), which is highly represented in RNA binding proteins (12,13). This domain mediates self-assembly and interactions with hnRNP complexes important for RNA processing activity (8,14,15), but at the same time, the CTD drives protein aggregation and toxicity (16–18). The CTD is characterized by an abundance of glutamine/asparagine residues, showing great similarity to prion domains in yeast proteins, such as that of the archetypal prion protein Sup35 (13,19). Significantly, almost all disease-associated TDP-43 mutations cluster in the CTD (5,20), strongly suggesting that these substitutions disrupt normal protein interactions and promote aggregate formation, driving the disease state.

The central mechanism in TDP-43 self-assembly and aggregation has been largely unexplored. TDP-43 aggregation assays using the full-length protein are encumbered by the extreme aggregation-prone characteristic of TDP-43, which makes production of pure soluble protein particularly challenging. Having recently established methods to generate soluble recombinant TDP-43 (21), we studied its aggregation to identify the factors that mediate and alter this process (e.g., ALS-associated mutations) and to gain insight into the structure of aggregates. We found that TDP-43 aggregates are formed through a biphasic process that initiates with oligomerization followed by aggregation into high molecular weight polymers. ALS-linked mutants potently affect aggregation by increasing the rate of assembly. In addition, we show that the aggregates derived from purified TDP-43 are capable of seeding intracellular aggregation following uptake. Our results support a model in which TDP-43 undergoes self-assembly into oligomeric complexes upon misfolding that act as templates for large aggregates. This process may be altered in disease conditions, such as in the presence of patient-linked mutations.

## RESULTS

### Initial TDP-43 oligomerization precedes aggregation into high molecular weight aggregates

We have successfully developed methods to generate soluble, full-length bacterial recombinant TDP-43 (rTDP-43) to characterize TDP-43 interactions (21) (**Fig. S1A**). We generated homogeneous and soluble TDP-43 fused to a SUMO tag at the N-terminus (21), which is cleaved off by the SUMO deconjugating protease Ulp1 (**Fig. S1B**). Using this protein, we analyzed TDP-43 self-assembly and aggregation. Prior to carrying out the aggregation assays, we ensured that purified protein preparations did not include pre-formed aggregates using highspeed ultracentrifugation and by routinely measuring protein activity in RNA binding assays (**Fig. S1C**). The binding affinity for the TDP-43-A(GU)_6_ RNA interaction was determined with our previously established fluorescence-based method (21). This accurately measures the apparent dissociation constant for A(GU)_6_ (typically *K_d,app_*=2.3 +/-0.7 nM) and also provides an estimate of the active protein concentration. By measuring these parameters and assuming that active protein concentration is equal to the concentration of soluble protein, we determined the purity of our protein preparations. Our aggregation assays consisted of analyzing rTDP-43 complexes before (day 0) and after brief shaking followed by incubation at room temperature (approximately 22ºC). To detect the assembled complexes, we adapted a method widely used for the characterization of yeast prion aggregates (22–24), based on the resolution of soluble and large aggregate complexes by semi-denaturing detergent agarose electrophoresis (SDD-AGE). SDD-AGE generates large pores allowing the detection of large protein complexes that are resistant to SDS detergent, typically ranging from 0.1-2% (24). This method has also been used to detect neurodegeneration-associated aggregates, such as tau (25), and in a limited number of studies to analyze TDP-43 pathology from patient tissue and cultured cells (26,27). We analyzed rTDP-43 aggregates formed over time and obtained the resolution of rTDP-43 monomers and large polymers, which increased in size (Fig. 1A). In addition to high molecular weight aggregates, we observed an oligomeric pattern of assembly at initial time points. These initial complexes (Figs. 1A, days 1-4, arrows) were replaced by larger, heterogeneous species at longer incubation times (Fig. 1A, days 8-15). The later day large complexes resembled aggregates formed by prion proteins and other protein aggregates analyzed with this method (22,25,28). Addition of polyethylene glycol (PEG), used as a crowding agent, accelerated the aggregation rate, increasing the formation of the larger complexes (**Fig. S2A**). Figure 1 shows complexes formed by SUMO-tagged TDP-43, and removal of SUMO had no significant effect in the size of the aggregates or in the formation of oligomers (**Fig. S2B**). In the absence of SUMO the assays were initiated immediately after removal of the tag as the cleaved TDP-43 could not be stored without significant protein loss. Therefore, we used SUMO-tagged protein in our experiments, unless noted.

**Figure 1.**
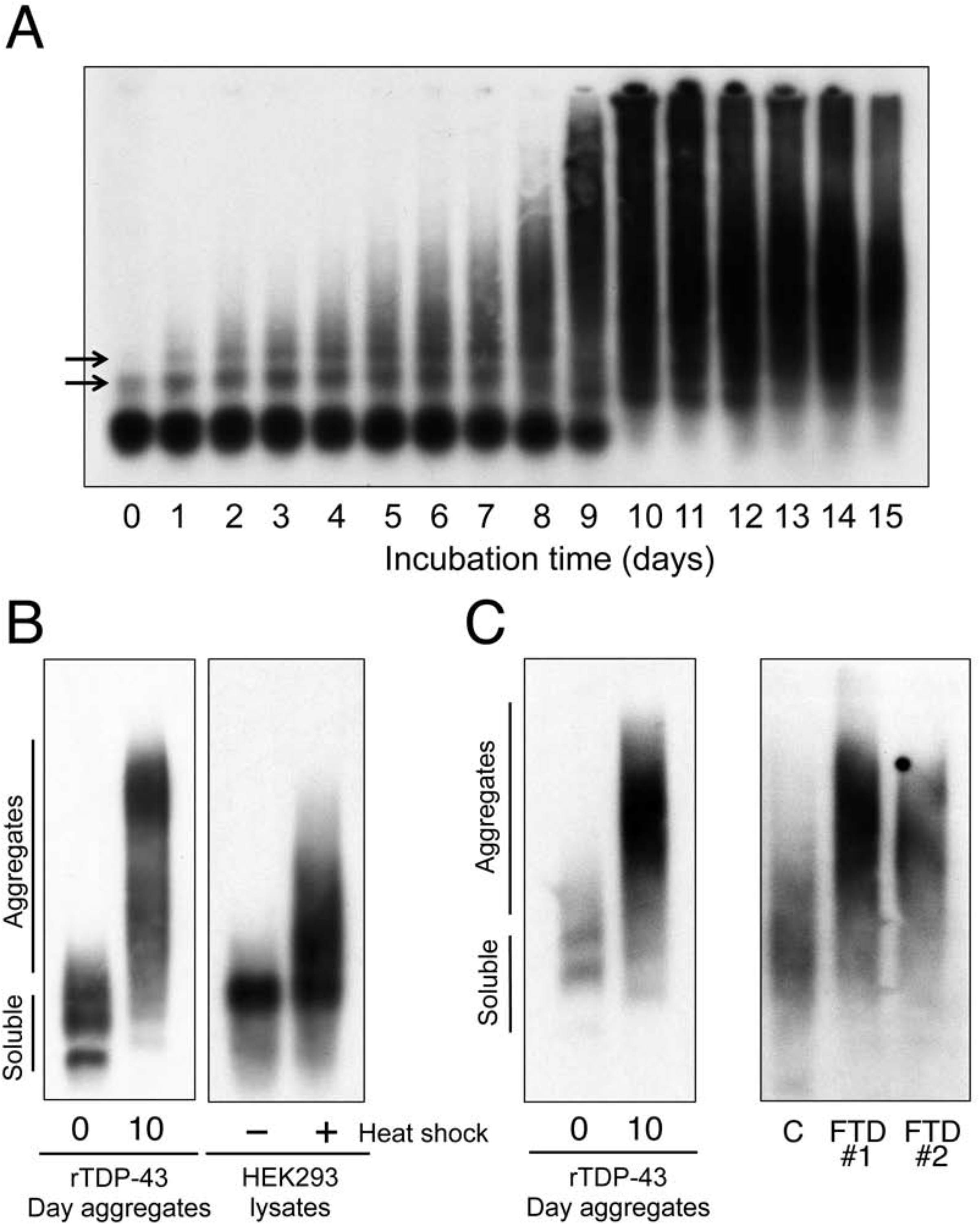
Detection of TDP-43 oligomers during the initial steps of aggregation. (A) Aggregation of full-length purified TDP-43 (rTDP-43) was triggered by brief shaking (2 μM in reaction buffer) and samples were analyzed as a function of time (days 0 to 15) by SDD-AGE/immunoblotting. Arrows point to the initial oligomeric species. (B) Comparison of day 0 and day 10 rTDP-43 aggregates to HEK293 cell lysates treated with heat shock (43ºC for 30 min) and non-treated control. (C) The sarkosyl insoluble fraction from FTD and control (C) brain samples analyzed by SDD-AGE. These were compared to Day 0 and day 10 rTDP-43 aggregates. rTDP-43 samples used as comparison (in B, C) were prepared in the same buffer and electrophoresis conditions as cell lysates and brain samples. All immunoblots were probed with TDP-43 antibody, shown are representative of > 3 individual experiments.

We compared aggregates from purified TDP-43 to cell-derived aggregates and insoluble TDP-43 fractions from FTD brain tissue samples to determine their physiological relevance and validate our methods. Lysates from human embryonic kidney (HEK293) cells treated with heat shock, which results in TDP-43 misfolding (21,29,30), showed high molecular weight complexes similar to the later day aggregates from rTDP-43 (Fig.1B). These were absent in non-treated control cell lysates. Next, we analyzed postmortem human brain tissue from control and FTD cases following sequential extraction of grey matter dissected from frontal cortex samples (Fig.1C). We used the sarkosyl insoluble fraction for our analyses as described in Methods (31). Our results showed that the levels of TDP-43 aggregates were reduced in the control sample compared with FTD, consistent with a previous report (26). In addition, the migration pattern of TDP-43 in control tissue was different from the disease cases. We included 0 and 10 day rTDP-43 aggregates in the same solution conditions as brain tissue for comparison (Fig.1C). This showed that migration of later day rTDP-43 aggregates was similar to TDP-43 complexes from FTD, but not from control tissue. Collectively, these results suggest that our methods could be used to detect and analyze aggregate species generated from the purified protein, which show conformational and biochemical similarities to cell and patient-derived TDP-43 inclusions.

We investigated whether TDP-43 complexes could be detected without triggering aggregation by shaking and found that 10 minute incubation of soluble protein at increasing temperatures enhanced the formation of oligomers as seen by SDD-AGE (Fig. 2A). The left panel of figure 2A shows aggregates formed at 0, 3, 5, and 10 days after shaking, for comparison. The TDP-43 complexes, which increase at higher temperatures are similar to the intermediate species in the aggregation assay. To estimate the oligomeric state of the early TDP-43 complexes we performed cross-linking experiments under reducing conditions (Fig. 2B). These complexes were analyzed by standard denaturing SDS-PAGE after the addition of b-ME (100 mM). As control, non-crosslinked samples were analyzed in the presence and absence of β-ME. Immunoblot analysis showed cross-linked complexes corresponding to tetramers at day 3, according to the estimated molecular weight, which were not detected in non-crosslinked samples independent of β-ME treatment. This suggests that oligomers form upon misfolding as the initial step of a two-stage process of aggregation, favoring a tetrameric complex. This early phase may act as the scaffold for larger aggregates.

**Figure 2.**
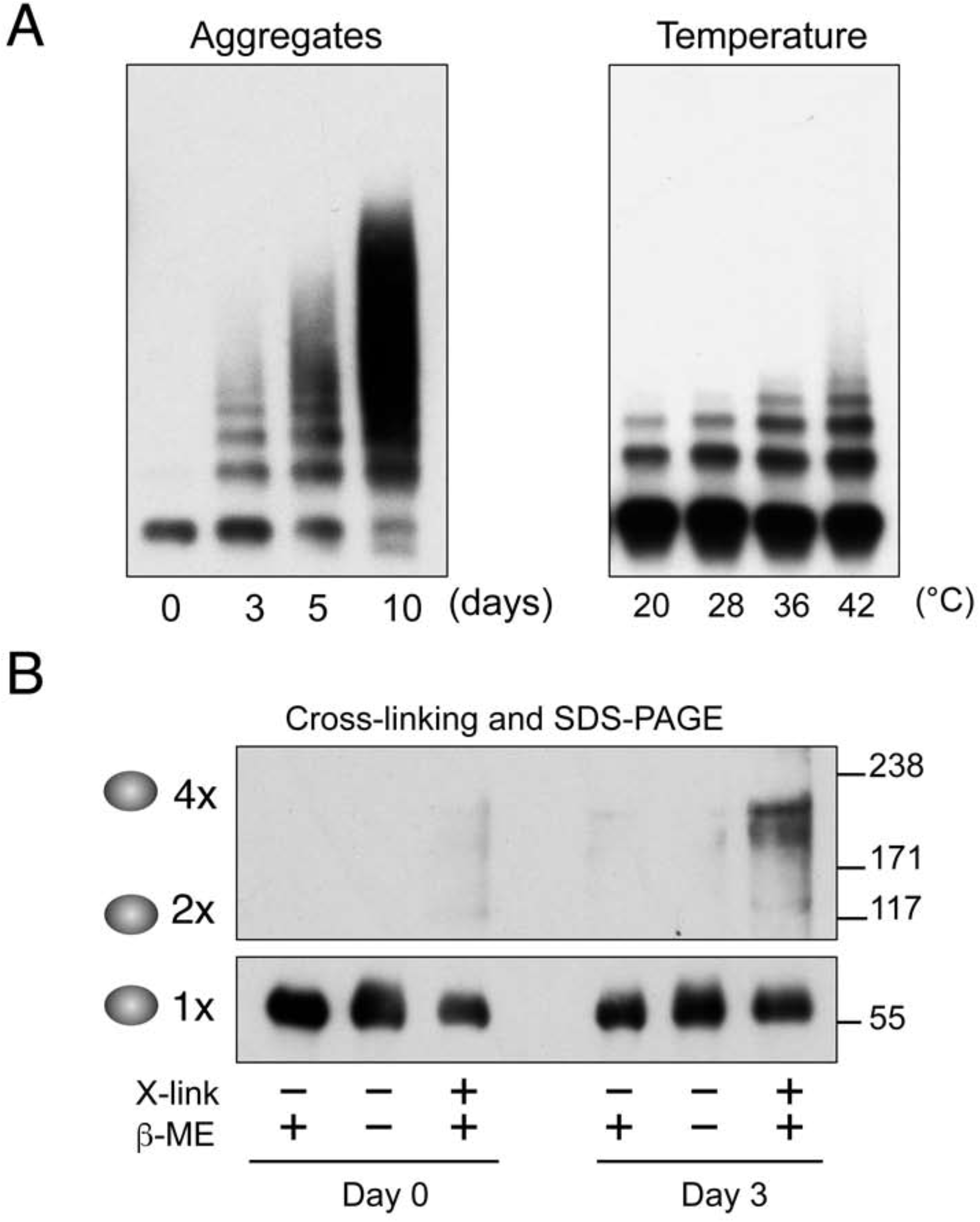
TDP-43 Oligomers favor tetramers and increase at higher temperatures. (A) SDD-AGE/immunoblotting analysis of aggregates formed at 0, 3, 5, and 10 days compared to TDP-43 complexes formed upon brief incubation at increasing temperatures without shaking (2 μM TDP-43 in reaction buffer for 10 minutes, 20 to 42ºC). (B) Days 0 and 3 TDP-43 complexes were chemically cross-linked (X-link) in the presence and absence of β- mercaptoethanol (250 mM β-ME). Samples were analyzed by SDS-PAGE and immunoblot. The estimated number of TDP-43 molecules cross-linked and the molecular weight markers (kDa) are shown. All blots are representative of > 3 individual experiments.

### Early TDP-43 complexes are not mediated by disulfide bonds

To further understand whether TDP-43 assembly was mediated by covalent interactions, we analyzed the sensitivity of the initial complexes and large aggregates to reducing agents. We used the reducing agent Tris(2-carboxyethyl)phosphine (TCEP) throughout rTDP-43 purification and during the aggregation assays. Moreover, we added supplementary TCEP on day 5 to ensure that the samples remained under reducing conditions (reducing, **Fig. S2C**). This was compared to a “nonreducing” assay, whereby aggregates were generated using rTDP-43 purified in the presence of β-ME, a less stable reducing agent compared to TCEP. No additional reducing agent was used in the aggregation assay (non-reducing, **Fig. S2C**). These non-reducing conditions generated large heterogeneous aggregates immediately, beginning at the initial time point (day 0). By comparing the results obtained under reducing and non-reducing conditions we found that the initial and intermediate aggregates were detected in the presence of TCEP, however, the larger aggregates decreased significantly under these conditions (i.e., day 10 in reducing and nonreducing samples). To further confirm our results, we treated aggregates formed at days 3, 7, and 14 with high concentrations of reducing agents (TCEP, β-ME and DTT, 250 mM) (**Fig. S2C**). This did not significantly affect the initial complexes, but greatly decreased formation of the largest aggregates at later time points. Based on these collective observations we propose that initial complexes that are detected as ordered molecular assemblies by SDD-AGE are partially detergent-resistant oligomers and are not mediated by covalent bonds. On the other hand, the higher molecular weight heterogeneous aggregates form through disulfide interactions. This is consistent with the effect of oxidative stress in promoting TDP-43 aggregation in cells and with the previous identification of intra and intermolecular TDP-43 Cys interactions in patient tissue (32). Our results and this previous work highlight the relevance of disulfide bond formation on TDP-43 pathology.

### Role of Liquid-liquid phase transitions in oligomer formation

RNA binding proteins associated with ALS/FTD, including TDP-43, FUS, TIA-1, Matrin 3 and hnRNP A1 undergo a process of condensation through liquid-liquid phase separation (LLPS) according to in vitro and in cell-based evidence (33). This activity mediates the formation of membraneless RNA-protein rich organelles in the nuclear and cytoplasmic cellular compartments (e.g., nucleoli and stress granules) (12,34-36). Liquid droplets assembled through LLPS “mature” over time into solid-like complexes such as fibrils. Importantly, disease-linked conditions are proposed to disrupt LLPS and RNA granule assembly, thereby promoting aggregation and neurotoxicity in ALS and FTD (37–39). Therefore, self-assembly of these ALS-FTD-associated proteins is central to their physiological function, but also plays a major role in aggregate formation. LLPS analysis of TDP-43 is still limited relative to the growing number of studies on FUS and other related proteins and this is likely due to differences in their phase separation properties (40). Therefore, the role of TDP-43 self-assembly or LLPS in aggregation and the mechanisms that connect these processes remain poorly characterized.

We asked whether the oligomeric complexes seen at the initial time points in the TDP-43 aggregation assay represent assemblies formed by LLPS. Microscope visualization of fluorescently labeled rTDP-43 in low salt (25-50 mM NaCl) showed immediate phase separation and formation of droplets similar to those recently reported by Maharana et al. (Fig. 3A). Droplet formation was salt-dependent as the number of droplets was dramatically reduced at higher salt concentrations (250 mM), consistent with previous studies (Fig. 3B) (11,36). Furthermore, phase separation was reversed by increasing the salt concentration to 250 mM, indicating that the observed complexes were not irreversible aggregates (data not shown). Interestingly, extending the incubation time of the sample beyond 30 minutes resulted in the growth of fibrillar structures as seen by fluorescence microscopy (Fig. 3A). We analyzed samples corresponding to those used for droplet formation at low and high salt and at longer incubation times by SDD-AGE (Fig. 3C). For control, we loaded samples from the aggregation assay days 0 and 10 (Figs.1A, 2A). We found no macromolecular complexes by SDD-AGE in droplet conditions (25-50 mM NaCl) or at high salt (250 mM). However, incubation of the sample for 2 hours, parallel to the conditions that resulted in liquid droplet maturation, showed an increase in the oligomeric species similar to the pattern detected at the initial time points of the aggregation assay (Fig. 3B). These observations suggest that maturation of droplets into assemblies with solid-like properties may be detected by SDD-AGE and provide further evidence that the detergent-resistant oligomers form as the initial steps of misfolding and aggregation.

**Figure 3.**
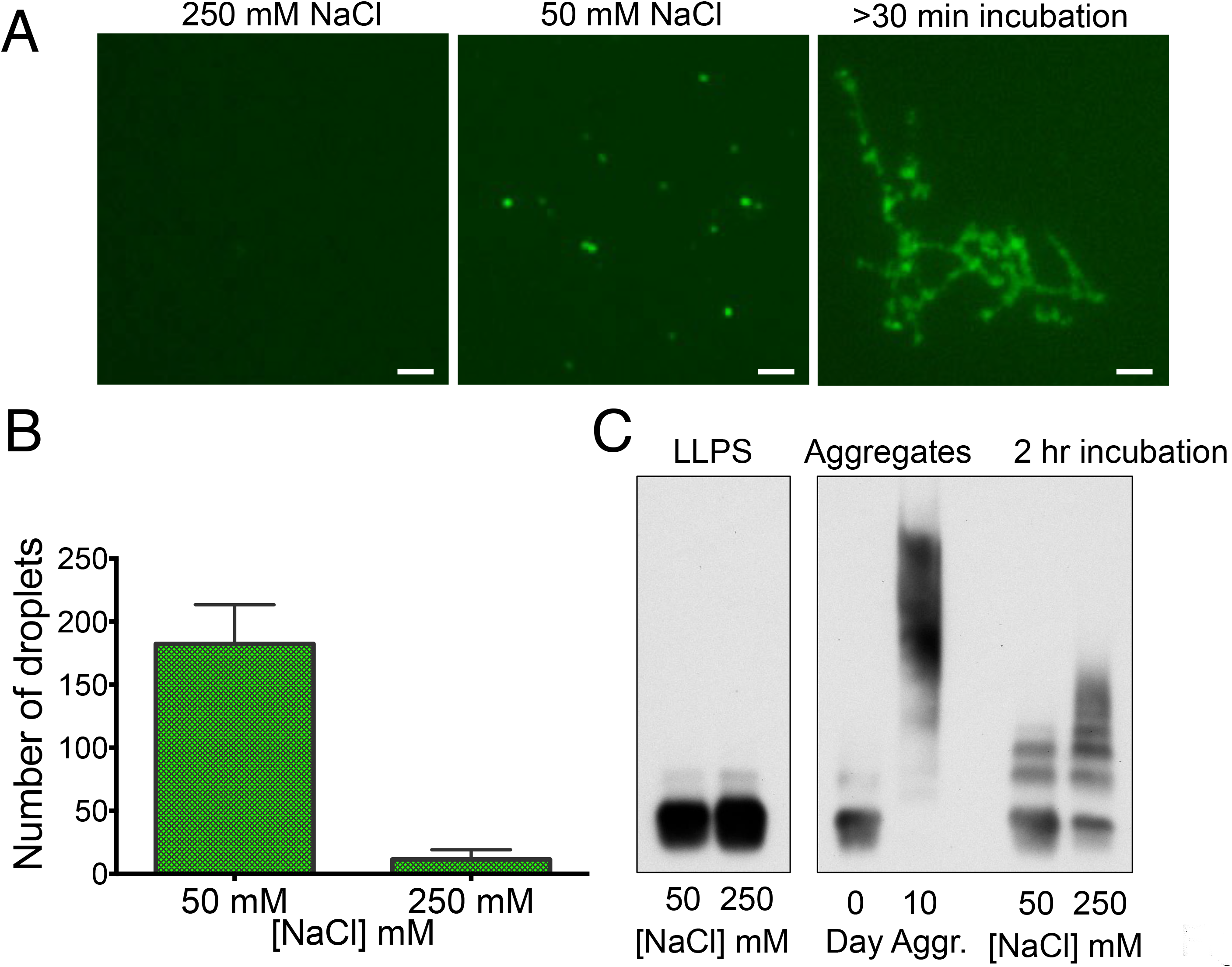
Maturation of TDP-43 liquid droplets is detected as oligomerization by SDD-AGE. (A) Representative microscope images of fluorescently labeled TDP-43 (0.5 μM) liquid-liquid phase separation (LLPS). Liquid droplets formed in low salt (50 mM NaCl), but not in high salt (250 mM NaCl). Incubation of samples for 30 minutes or longer at 22ºC resulted in fibril-like structures. Scale bar, 2 μm. (B) The number of droplets formed at 50 and 250 mM NaCl were quantified by ImageJ (n = 3, s.e.m.). (C) Samples in low and high salt conditions analyzed by SDD-AGE/immunoblot and compared to 0 and 10 day aggregates formed as in Fig.1A. LLPS samples at low and high salt analyzed after prolonged incubation (2 hours). Microscope images and blots are representative of > 3 individual experiments.

### Contribution of TDP-43 domains on protein assembly

To determine how each of the modular domains contributes to the two stages of assembly, we compared complex formation and aggregation of rTDP-43 deletion mutants (Fig. 4A). SDD-AGE analysis of the complexes showed that removal of the amino-terminal domain (ΔN) decreased the formation of the initial oligomers and increased the formation of the large aggregates relative to the full-length protein (Fig. 4B). Conversely, deletion of the C-terminal low complexity region (ΔC) maintained the pattern of oligomerization and reduced the levels of large aggregates, consistent with its prionlike function. Removal of both N-and C-terminal domains (RRM1-2) triggered the formation of multiple oligomeric complexes starting at day 0 with little accumulation of high molecular weight species even after prolonged growth time. These results were supported by solubility assays in which protein samples (days 0 and 5 of the aggregation assay) were fractionated into RIPA-soluble and-insoluble fractions (Fig. 4C). The ratio of Urea-soluble to RIPA-soluble protein was similar for all constructs at the initial time point (day 0). As expected, the later time point showed a significant increase of insoluble protein for full-length and ΔN (day 5) (Fig. 4C,D). Deletion of the C-terminal low complexity domain reduced insoluble aggregates 6-fold compared to full-length TDP-43. The RRM1-2 fragment showed the highest solubility with little increase in the insoluble fraction over time (Fig. 4C,D). Collectively, our results indicate a strong contribution of the amino-terminal and RRM domains to TDP-43 self-assembly into oligomeric complexes. On the other hand, the C-tail plays a critical role in the formation of large aggregates consistent with the strong aggregation-prone characteristic of this domain (16–18). These observations also suggest that the N-terminal domain is important for TDP-43 oligomerization in the presence of the C-terminal tail, in the context of full-length protein. This model is in agreement with studies showing that oligomerization mediated by the N-terminal domain counteracts aggregate formation in cells, suggesting that this interaction controls the assembly of the full-length protein (9,11). However, our results showed that in the absence of the disordered C-tail, the RRM domains are sufficient to promote oligomerization without the requirement of the N-terminal domain. Future studies will further map the regions of interaction that mediate oligomerization to determine whether preventing oligomerization through RRM1-2 interactions affects aggregation.

**Figure 4.**
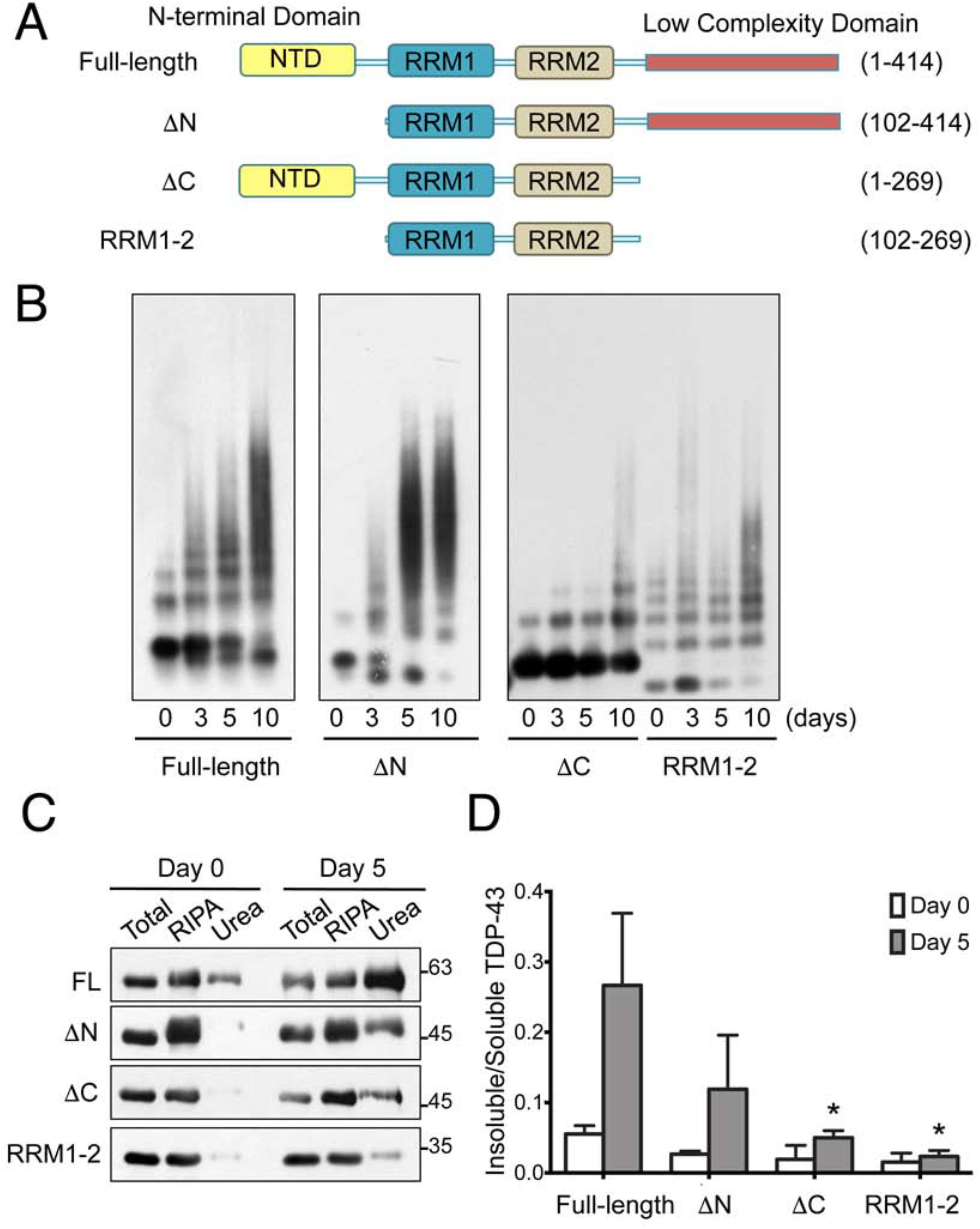
Role of TDP-43 protein domains in oligomerization and aggregate formation. (A) Schematic representation of TDP-43 deletion constructs devoid of N-terminus (ΔN), C-terminus (ΔC), or both (RRM1-2) consisting of the amino acid residues shown. (B) Aggregates formed by these protein fragments were analyzed by SDD-AGE/immunoblotting. Samples from Days 0, 3, 5, and 10 were selected for comparison with full-length TDP-43. (C) SDS-PAGE and immunoblotting of full-length rTDP-43 (FL) and deletion fragments fractionated into RIPA soluble and insoluble fractions. Insoluble pellets were resuspended in urea at days 0 and 5 of the aggregation assay. Equal volumes of starting lysate (Total) and RIPA-soluble fraction were loaded. Urea soluble pellet represents 5-fold concentrated sample relative to RIPA-soluble fraction. (D) RIPA and urea soluble TDP-43 was quantified to compare the levels aggregation at the initial time point (day 0) and day 5 of the aggregation assay. Band intensity was quantified by ImageJ (n = 3, s.d.). The statistical significance (*P< 0.025) comparing deletion fragments to full-length rTDP-43. All blots are representative of > 3 individual experiments.

### rTDP-43 oligomers are structuraly diferent from late-stage aggregates

We performed atomic force microscopy (AFM) and immuno-electron microscopy (EM) to further investigate the structural changes associated with the different stages of TDP-43 aggregation seen by SDD-AGE. Using these techniques, we studied both full-length TDP-43 and the RRM1-2 fragment to compare the structure of large aggregates versus oligomeric patterns given that RRM1-2 complexes resembled early day full-length oligomers. AFM experiments showed that RRM1-2 does not form visible particles at day 0, whereas the full-length protein revealed small round oligomeric structures without incubation (Fig. 5A). After one day, the full-length formed larger density oligomers compared to RRM1-2. Nascent fibrillar structures of full-length rTDP-43 were observed after two days incubation, but were absent in RRM1-2 (Fig. 5A). Similarly, EM analysis showed fibrils of full-length TDP-43 at day 2, which grew in size after 7 days of incubation (Fig. 5B). This is consistent with previously obtained complexes formed by full-length TDP-43 and C-terminal domain protein fragments (27,41). In contrast, RRM1-2 at day 7 formed large oligomers under the same conditions. Only after prolonged incubation (day 10), RRM1-2 formed large aggregates with fibrillar substructure as seen with full-length TDP-43. The lack of fibrillar structures up to day 7 in the RRM1-2 complexes were consistent with the oligomeric pattern seen by SDD-AGE. Binding of anti-TDP-43 immuno-gold particles confirmed that the assemblies were formed by the respective proteins. Collectively, these results strongly suggest that the initial assembly states during TDP-43 aggregation, which we detected as an ordered oligomeric pattern by SDD-AGE, are structurally different from later day aggregates. The larger aggregates form fibril-like structures similar to those previously observed for full-length TDP-43 using similar microscopy methods (16,41).

**Figure 5.**
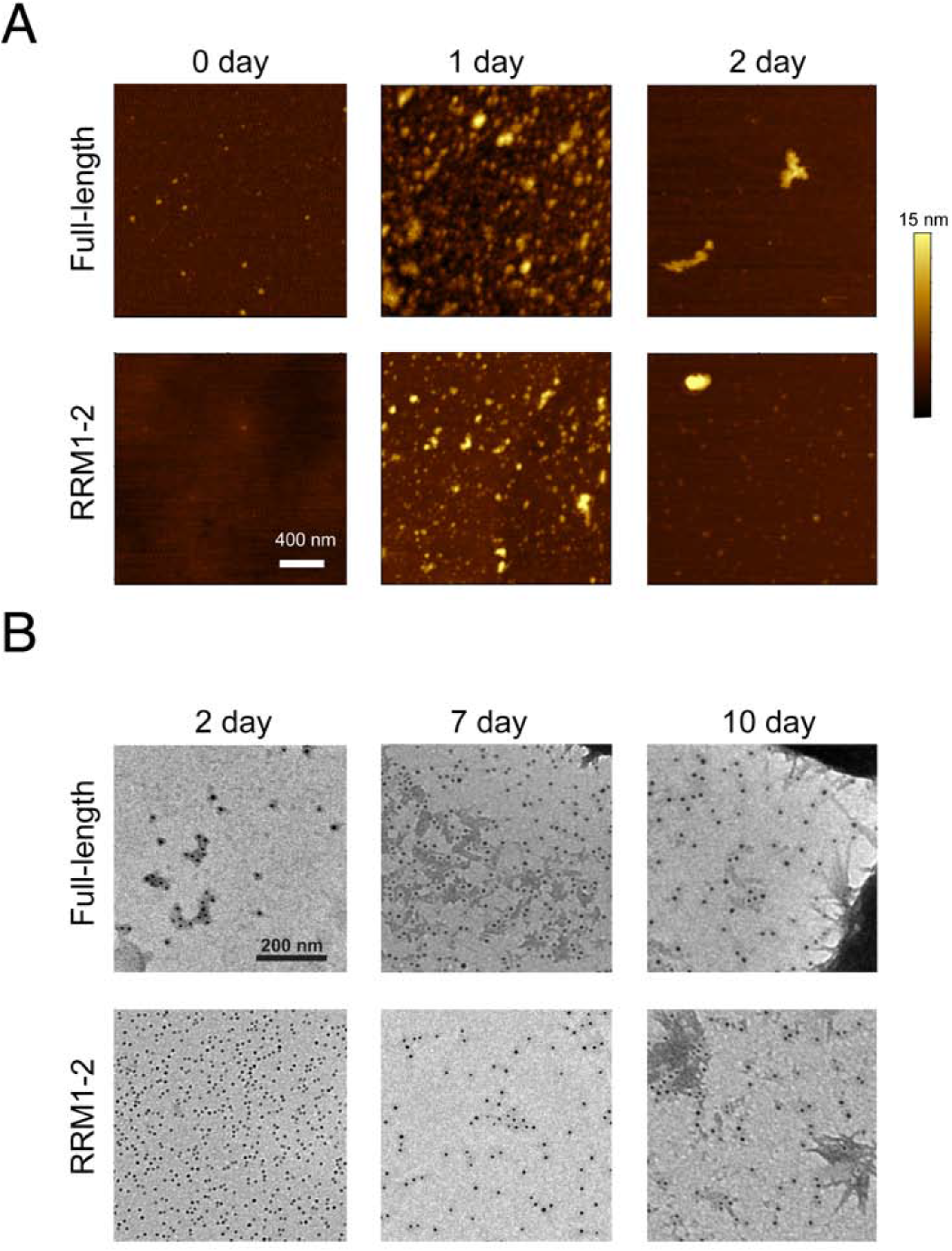
Oligomers are conformationally distinct from the large late-stage aggregates. Structural analyses of full length TDP-43 and RNA recognition motif fragment (RRM1-2) to analyze soluble, early (0, 1, 2 day) and late assembly samples (7 and 10 day). (A) Representative atomic force microscopy image of the complexes at each time point. Scale bar, 400 nm. Z color bar, 0-15 nm. (B) Immuno-electron microscopy of the aggregates using TDP-43 antibody coated particles. Scale bar, 200 nm. Full-length TDP-43 formed fibrillar structures faster than RRM1-2 fragments.

### Disease-linked mutations increase the assembly and aggregation of TDP-43 in vitro

The mechanism by which disease-associated mutations promote neurodegeneration remains unclear. To test whether our assay may be used to determine and measure changes in the aggregation process caused by these mutations, we selected two substitutions: A315T and M337V (Fig. 6A). Both mutations are highly associated with disease and previously reported to increase aggregation in vitro and promote neurotoxicity in animal models (5,27,42,43). We performed aggregation assays with A315T and M337V and analyzed the complexes formed over time by SDD-AGE (Fig. 6B). We observed that both mutations potently impacted the rate of aggregation as seen by significantly accelerated oligomer formation at the earliest time point (day 0), which was mostly absent in wild-type. The oligomer pattern typical of days 3 and 5 in wild-type TDP-43 was substituted by the larger aggregates in the mutants, particularly in the case of M337V. Quantifying aggregation rates from the loss of monomer protein at different time points showed that while wild-type monomer decreased linearly as a function of time, A315T and M337V monomers decreased at a significantly faster exponential rate (Fig. 6C). This, particularly the effect of M337V, is in strong agreement with previous observations that these substitutions increase amyloid formation of the isolated disordered C-tail (44,45) and full-length protein (16) and disrupt phase separation in cell models (46–48). Our results on the macromolecular assembly of A315T and M337V may reflect distinct structures formed in the presence of the substitutions that have faster rates of complex association. Alternatively, these results may suggest that disease mutations decrease the dissociation of TDP-43 molecules making more stable complexes that result in stable fibrils or aggregates.

**Figure 6.**
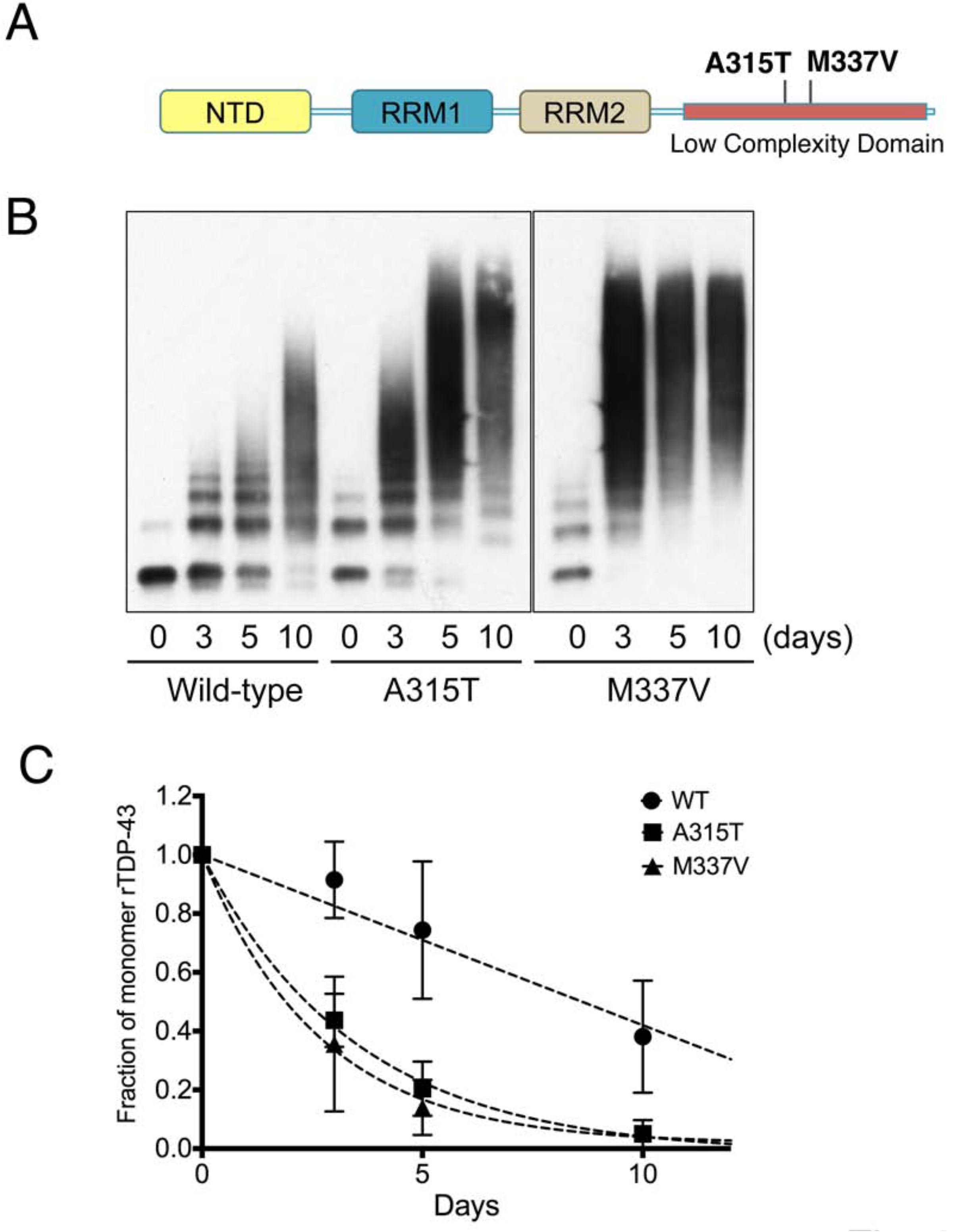
ALS mutations accelerate the rate of TDP-43 aggregation into high molecular weight aggregates. ALS-linked mutations A315T and M337V in the C-terminal domain of TDP-43 (A) were analyzed by SDD-AGE/immunoblot (B). Samples were taken at initial time 0 and days 3, 5, and 10 of the aggregation assay. Blots are representative of > 3 individual experiments. (C) Monomer, corresponding to the fastest migrating band, was quantified by ImageJ at the different time points for wild-type, A315T and M337V. Curves were generated by fitting data points to an exponential formula in GraphPad Prism (n = 3, s.e.m.).

### rTDP-43 aggregates seed and propagate intracellularly and this is enhanced by ALS mutations

Propagation of aggregates through prion-like transcellular transmission has been documented for a number of proteins associated with neurodegeneration, including TDP-43 (49,50). We sought to investigate whether our rTDP-43 aggregates were capable of seeding aggregation in cells. For this, we generated a stable cell line to detect cytoplasmic aggregates in a reproducible and sensitive assay. A single copy of mCherry-tagged TDP-43 was stably integrated in HEK293 cells for tetracycline-induced expression (HEK-TDP-43^NLS^) (**Fig. S3**). This allowed easy visualization of TDP-43 and prevented high and non-homogeneous levels of transgene overexpression that could disrupt protein homeostasis. This system offers an advantage to studying TDP-43 aggregation relative to transient transfection, which often causes overexpression of the protein. While TDP-43 shuttles between nucleus and cytoplasm (51), it is predominantly nuclear. We expressed full-length TDP-43^NLS^ harboring alanine substitutions to disrupt the nuclear localization signal (NLS) and increase cytoplasmic TDP-43, as previously observed (52,53) (**Fig. S3A,B**). Expression of a similar TDP-43 NLS-mutant causes neurotoxicity accompanied by pathological inclusions in brain and spinal cord in mice (54). Our HEK-TDP-43^NLS^ cells expressed soluble TDP-43^NLS^ under control conditions, but could be induced to form cytoplasmic aggregates readily detected by microscopy after addition of proteotoxic agents (e.g., MG132 and sodium arsenite) (**Fig. S3B,E**). We confirmed that the cytoplasmic inclusions triggered by proteotoxic stress were recognized by established markers of TDP-43 pathology, i.e., phosphorylation at Ser409/410 (55,56) (**Fig. S3C,D**).

To test whether the aggregates generated from rTDP-43 in vitro could be internalized in HEK-TDP-43^NLS^ and nucleate intracellular aggregation, we fluorescently labeled rTDP-43 with Oregon Green before initiating the aggregation assays. Aggregates from day 5 (Fig. 1A) were added to cultured HEK-TDP-43^NLS^. Cells were trypsinized and replated 24 hours post-transfection to remove rTDP-43 from the media and from association with the extracellular membrane. We observed internalization of labeled rTDP-43 48 hours post transfection (Fig. 7A). Aggregate formation of mCherry-TDP-43^NLS^ was monitored for several days. We observed a significant increase in cellular TDP-43 aggregation after 6 days of treatment, as seen by formation of large mCherry cytoplasmic foci (Fig. 7A). Importantly, these colocalized with the Oregon Green-labeled rTDP-43 signal. Cells treated with rTDP-43 aggregates showed a significant 2-fold increase of cytoplasmic aggregates compared to those treated with soluble rTDP-43 or no protein control (Fig. 7B), suggesting that aggregates derived from purified TDP-43 may propagate misfolding once internalized in cells. These aggregates were also positive for pSer409/410 (**Fig. S4A, B**), suggesting that the inclusions share structural and mechanistic similarities with those formed in disease. Previous studies showed TDP-43 seeding in cells, however these assays used recombinant TDP-43 protein fragments and the resulting intracellular aggregates were not detected by pSer409/410 (57).

**Figure 7.**
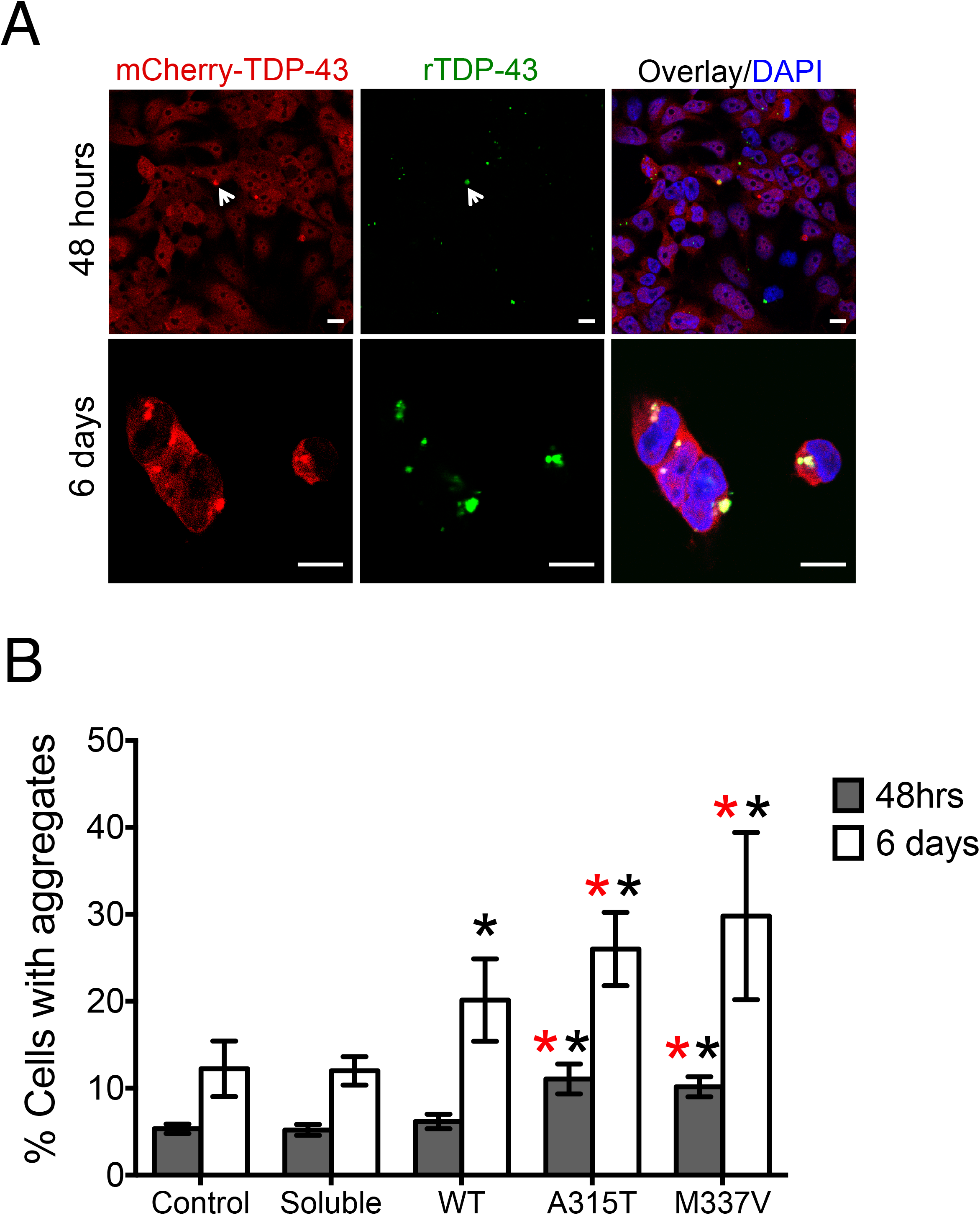
rTDP-43 aggregates propagate intracellularly and more efficiently with disease-linked mutations. (A) Oregon Green-labeled rTDP-43 aggregates, collected briefly after shaking for 30 min, transfected in mCherry-TDP-43 expressing cells (HEK-TDP-43^NLS^). Cells were trypsinized 24 hours post-transfection and rTDP-43 aggregate internalization was monitored 48 hours post-transfection (top panel). Arrowhead points to colocalization of Green aggregates with mCherry-TDP-43 aggregates in the cytoplasm. Bottom panel, mCherry-tagged TDP-43^NLS^ cellular aggregates visualized by microscopy at higher magnification after 6 days of treatment. Overlay shows additional merging with DAPI. Scale bar=10μm. (B) The percent of HEK-TDP-43^NLS^ cells showing cytoplasmic aggregates (mCherry-positive) was quantified after 48 hours and 6 days of transfection with rTDP-43, including wild-type rTDP-43 aggregates (WT) and ALS mutant rTDP-43 aggregates (A315T and M337V). No protein and soluble wild-type rTDP-43 transfection were used as control. Statistical significance was calculated for control vs. WT, A315T and M337V at 48 hrs and 6 days *; WT vs. A315T and M337V at 48 hrs and 6 days *(>200 cells/sample counted in n > 3, s.d., GraphPad Prism two-tailed t-test analysis * * P≤ 0.04).

Next, we analyzed whether the ALS/FTD-linked mutations altered aggregate formation in cells as seen in our in vitro assays. Transfection of HEK-TDP-43^NLS^ with A315T and M33ZV rTDP-43 aggregates showed a significant 2-fold increase in mCherry-positive aggregates already at 48 hours (Fig. 7B), whereas wild-type rTDP-43 aggregates did not show a significant difference compared to control at this time point. The effect of the disease-associated mutations was even greater after 6 days, showing a 3-fold increase in mCherry-TDP-43^NLS^ aggregates compared to control (Fig. 7B). A315T and M33ZV also caused significantly greater intracellular aggregation compared to wild-type, 50 and 67% increase, respectively, 6 days post-transfection. Of note, cellular internalization of rTDP-43 aggregates was similar for wild-type and mutant, approximately 40% of cells showed cytoplasmic labeled rTDP-43. These results strongly suggest that the mutants do not alter cellular uptake, but are more efficient at seeding relative to wild-type. This is in agreement with the increased rate of large aggregate assembly seen by SDD-AGE in vitro (Fig. 6B).

## DISCUSSION

Our studies shed new light into the molecular steps that lead to TDP-43 aggregation and highlight methods to study the factors that control TDP-43 proteostasis and/or trigger protein misfolding. We also report a previously unappreciated role for disease-associated mutations on the aggregation mechanisms, providing important clues on how these mutations lead to pathology.

The use of SDD-AGE to analyze rTDP-43 aggregates provided the first evidence of a defined oligomeric pattern that precedes formation of TDP-43 high molecular weight aggregates (Fig. 1B, 2A). These intermediates may not be observed by other methods used to monitor aggregation, such as turbidity and other spectroscopic techniques. We confirmed the differences in the conformation and assembly state of early and late stage aggregates by AFM/immuno-EM analysis (Fig. 5). Interestingly, our findings suggest that the oligomers detected by SDD-AGE are key intermediates during the transition from complexes formed through liquid phase separation into irreversible aggregates (Fig. 3). This is consistent with the model that TDP-43 liquid-liquid phase separation may lead to aggregation over time, as suggested by studies of similar RNA binding proteins linked to ALS/FTD(37,39). This mechanism resembles the aggregation process of other neurotoxic proteins, such as tau, α- synuclein and amyloid-beta where these intermediate products have been proposed to be the toxic seeds that transmit cell damage and pathology (58–64). Future experiments will examine whether the TDP-43 oligomers at the initial aggregation time points are also more toxic to cells and/or propagate more efficiently than large aggregates in animal models.

The mechanisms and structural determinants of TDP-43 oligomerization have not been widely reported, although strong evidence supports TDP-43 physiological oligomerization in cells and in brain tissue (9,27,65). The initial complexes we observe in our aggregation assays represent at least partially misfolded species that act as precursors of protein aggregation. These are likely to be different from the soluble assemblies formed in cells, which would be expected to disassemble under the semi-denaturing conditions used for SDD-AGE analysis. Our SDD-AGE results showing that the size and levels of oligomeric complexes increases at higher temperatures further suggest that these form as the protein undergoes misfolding (Fig. 2A). Based on our findings, TDP-43 shows key similarities with the canonical yeast prion protein Sup35. This protein was recently found to assemble into functional, soluble complexes through liquid-liquid phase separation in vitro and in yeast models (66), and also form partially SDS-resistant oligomers that promote prion propagation (67).

Our observation that aggregates generated from the purified protein seed cellular cytoplasmic inclusions (Fig. 6) are consistent with previous reports and our own observations (data not shown) that TDP-43 aggregates from disease tissue and isolated TDP-43 C-terminal fragments promote cellular TDP-43 aggregation (50,68,69). Moreover, these results show that rTDP-43 aggregates are relevant to cell and patient-derived inclusions and suggest that, similar to tau and α-synuclein, TDP-43 aggregate seeds template and propagate transcellularly (49). We also provide strong evidence that enhanced TDP-43 aggregation rates translate into greater cellular transmission of aggregates, as seen with the ALS-linked mutations A315T and M337V. By studying the aggregation and propagation of A315T and M337V we propose that pathogenesis is strongly influenced by factors that increase the assembly and/or alter the dissociation rates. Changes in the aggregation rate may be due to structurally distinct complexes associated with the mutations. However, our observations that A315T and M337V aggregates nucleate cellular TDP-43 suggest that mutant complex structure and assembly pathways are sufficiently similar to wild-type to act as seeds for aggregation. These findings open the avenue for future studies aimed at addressing whether the mechanisms observed with A315T and M337V apply to other TDP-43 disease-associated mutations. This will increase our understanding of the molecular basis of ALS-FTD pathogenesis. Importantly, studies on the mechanisms by which these mutations alter structural and kinetic parameters of TDP-43 self-assembly will shed light on the regulation of the equilibrium between soluble complexes and irreversible aggregates.

## EXPERIMENTAL PROCEDURES

All reagents are from Sigma-Aldrich unless otherwise specified.

### Plasmid construction

Construction of the TDP-43 vector for bacterial expression (SUMO-TDP43) was previously described (21). Oligonucleotides used for mutagenesis and cloning are described in Supplementary Table 1. Deletion constructs for bacterial expression were made by standard PCR and cloning between BamHI and SacI restriction sites in pET28b/His-SUMO (70). Single site substitutions including modification of the TDP-43 nuclear localization sequence, were made by site-directed mutagenesis using the QuickChange Site-Directed Mutagenesis Kit (Agilent Technologies) as described (15). The template used for mutagenesis of the mammalian expression vector, HA-mCherry-TDP-43, was made by cloning hemaggluttinin peptide-mCherry cDNA into pCDNA5/FRT upstream of TDP-43 cDNA (NM_007375.3). TDP-43 was cloned between BamHI and NotI sites.

### Recombinant TDP-43 production and aggregation assays

Recombinant TDP-43 (rTDP-43) was generated in *E.coli* and purified as previously described (21). His-tagged recombinant Ulp1 protease was used to remove SUMO. Briefly, rTDP-43 in the expression lysate was bound to Ni-NTA agarose and washed with Wash Buffer 1 (50 mM Tris, pH 8.0, 500 mM NaCl, 10% Glycerol, 10% Sucrose, 1 mM TCEP), Wash Buffer 2 (50 mM Tris, pH 8.0, 500 mM NaCl, 10% Glycerol, 10% Sucrose, 50 mM Ultrol Grade imidazole, pH 8.0, 1 mM TCEP), and finally washed again with Wash Buffer 1 to remove imidazole. Ulp1 was added to the protein-bound beads at an approximate 1:1 molar ratio and incubated for 30 min at 22°C. The supernatant containing cleaved TDP-43 was collected. For the aggregation assays rTDP-43 was ultracentrifuged in a Beckman Coulter Optima TLX Ultracentrifuge using a TLA55 rotor at 40,000 rpm for 30 minutes at 4°C to remove any pre-existing aggregates. Soluble protein concentration was measured by nanodrop and diluted to 2 μM in the reaction buffer: 50 mM Tris, pH 8.0, 250 mM NaCl, 5% glycerol, 5% sucrose, 150 mM Ultrol Grade imidazole, pH 8.0. rTDP-43 aggregation was started by shaking at 1,000 rpm, 22°C, for 30 minutes with an Eppendorf ThermoMixer C. Samples were incubated at 22°C and collected for analysis by adding equal volume of 2X SDD-AGE sample buffer (80 mM Tris-HCl, pH 6.8, 10% glycerol, 1% SDS, 0.2% bromophenol blue) and incubated at 22°C for 10 min prior to loading on SDD-AGE. Analysis of rTDP-43 aggregates was preformed by SDD-AGE as previously described (24,28), on a horizontal 1.5% agarose gel electrophoresis in 20 mM Tris, 200 mM glycine and 0.1% SDS. Proteins were transferred onto PVDF membrane (Amersham Hybond 0.45 mm PVDF, GE Healthcare) in a modified Mini Trans-Blot Cell (Bio-Rad) at 4°C. Protein was detected with traditional immunoblotting.

### Sequential extraction of TDP-43 from postmortem human brain tissue

Samples of frozen postmortem human brain tissue from Control and FTD de-identified cases were obtained from the Cognitive Neurology and AD Center (CNADC) at Northwestern University. Grey matter was dissected from frontal cortex samples using a scalpel, while maintaining the tissue in a frozen state. Extraction methods were adapted from (31). Samples of dissected tissue (approximately 100 mg) were homogenized in 5 mL/g Tris-sucrose buffer (25 mM Tris-HCl, pH 7.4, 5 mM EDTA, 1 mM DTT, 10% sucrose, protease inhibitor (PI) Cocktail) with Kontes Dounce Tissue Grinders (Kimble KT885300-0002). The homogenate was spun at 18,000 x g (average RCF) for 30 min at 4 °C. The resulting pellet was next extracted in 5 mL/g Triton X (TX) buffer (Tris Sucrose buffer, 1% Triton X-100, 0.5 M NaCl) and centrifuged at 135,000xg (RCF average) for 30 min at 4 °C. To float and remove myelin, the pellet was further extracted in “TX-30% Sucrose” buffer (25 mM Tris-HCl, pH 7.4, 5 mM EDTA, 1 mM DTT, 10% sucrose, 1% TX-100, 0. 5 M NaCl, 30% sucrose, PI Cocktail) and centrifuged at 135,000xg (RCF average) for 30 min at 4 °C. Finally, the pellet was homogenized in 5 mL/g Sarkosyl buffer (25 mM Tris, pH 7.4, 5 mM EDTA, 1 mM DTT, 10% sucrose, 1x PI Cocktail, 1% N-Lauroylsarcosine sodium salt, 0.5 M NaCl) and incubated at room temperature for 1h with shaking at 750 rpm on Advanced Vortex Mixer (VWR 89399-884). The Sarkosyl extracted homogenate was centrifuged at 135,000xg (RCF average) for 30 min at 4 °C. The resulting pellet was resuspended in 100 mL of RIPA buffer (50 mM Tris, pH 8.0, 150 mM NaCl, 1% nonylphenyl-polyethylene glycol (NP-40), 0.5% SDC, 0.1% SDS, 5 mM EDTA, PI cocktail, PhosStop (Roche)), sonicated with a nanoruptor (Diagenode) 30 sec ON, 30 sec OFF for 10 cycles. SDD-AGE analysis was carried out with 25 mL of this sample, representing 12.5% of the initial sample.

### Liquid-liquid phase separation assays

Oregon green 488 labeled rTDP-43 was mixed with unlabeled protein at a 1:10 ratio at 0.5 μM in 18 mM MES, pH 7.0, 5 mM Tris, 50 mM NaCl, 1% Glycerol, 1% Sucrose, 30 mM Imidazole. Samples were visualized by fluorescence microscopy immediately or incubated at 22°C for 30-120mins prior to analysis or SDD-AGE. Reversal of phase separation was achieved by raising the NaCl to 250mM immediately after droplet formation.

### Insoluble/soluble fractionation of rTDP-43 proteir 7s

Samples were collected after ultracentrifugation as above (Day 0) or shaken for 30 min and collected after 5 days of incubation at 22°C (Day 5). Samples were sonicated in a Bioruptor Pico (Diagenode) (30sec on / 30sec off, 10 cycles) and ultracentrifuged at 40,000 rpm for 30 min at 4°C and soluble fraction was obtained. The insoluble pellet was washed with RIPA-50 buffer (50 mM Tris, pH 8.0, 150 mM NaCl, 1% NP-40, 10 mM (ethylenedinitrilo)tetraacetic acid (EDTA), 0.5% sodium deoxycholate (SDC), and 0.1% SDS), sonicated and ultracentrifuged. The remaining insoluble pellet was resuspended in urea buffer (30 mM Tris, pH 8.8, 7 M urea, 2 M thiourea, and 4% 3-[(3-cholamidopropyl)dimethylammonio]-1-propanesulfonate hydrate (CHAPS)).

### Crosslinking and labeling rTDP-43

For crosslinking analysis, rTDP-43 (0.5 or 1 μM) was incubated with 0.25 mM disuccinimidyl suberate (DSS, Thermo Scientific) for 1 h at 22°C. The reaction was quenched by addition of Tris, pH 8.0 to 50 mM final concentration and incubated for 15 minutes at 22°C. rTDP-43 was labeled with Oregon Green 488 maleimide (ThermoFisher Scientific) according to manufacturer protocol. Labeled protein was purified with ZebaSpin columns (ThermoFisher Scientific) and used to label aggregates. Aggregation or LLPS assays included a 1:10 ratio mixture of labeled to non-labeled rTDP-43.

### Atomic force and Immuno-electron microscopy

Aliquots of rTDP-43 aggregation time points (10 μL) were placed on a clean, freshly cleaved grade V-1 mica (Cat#: 01792-AB, Structure Probe, Inc., USA). After 10 min, the solvent was wicked off by filter paper and the mica was washed 6 times with 20 μL of water to remove salts and buffer from the sample. Samples were dried overnight, and AFM images were acquired in tapping mode on a Veeco Dimension 3100 machine (Bruker) with Bruker FESP tips. AFM images were analyzed using the Gwyddion SPM data visualization tool. Carbon films on 200-mesh copper grids (Ted Pella) were incubated with 5 μL of sample in the dark side of the grid. After 10 min the sample was wicked off from the grid and was incubated with 10 μL, 1% BSA in PBS to block any nonspecific binding. The grid was incubated with primary rabbit TDP-43 antibody 1:100 diluted in PBS solution containing 0.1% BSA for 45 min. The grid was then washed by 7 drops of PBS buffer. The grid was incubated with secondary anti-rabbit IgG antibody conjugated to 5 nm diameter gold nanoparticles (Sigma Aldrich) diluted 1:20 in PBS buffer containing 0.1% BSA. 45 min later the grid was washed by 7 drops of PBS followed by 7 drops of water. Finally, the grid was stained by 2% uranyl acetate solution for 2 minutes and air dried before collecting the images on a FEI Transmission Electron Microscope.

### Cell culture

Cells were grown in growth media-Dulbecco’s Modified Eagle’s medium, 4,500 mg/L glucose, L-glutamine, and sodium bicarbonate and supplemented with filtered fetal bovine serum at 10%. Cells were incubated at 37°C, 5% CO_2_. Stable human embryonic kidney cells expressing TDP-43^NLS^ (HEK-TDP-43^NLS^) upon induction with tetracycline were generated from Flp-In™T-REX™293 cells (ThermoFisher) according to manufacturer instructions. Cells were grown in the presence of hygromycin (50 μg/mL) and transgene expression was induced with 1 μg/mL tetracycline.

### TDP-43 aggregate cellular popagaion

HEK-TDP-43^NLS^ were plated and induced with 1 μg/mL tetracycline for 16 h to reach 80-90% confluency. Recombinant TDP-43, Day 5 aggregates were transfected with Pierce™ Protein Transfection Reagent according to manufacturer instructions at a final protein concentration 0.1 μM in the culture media. As controls, soluble rTDP-43 or transfection reagent only were used. Media was changed 6 h post transfection. After 24 h, cells were trypsinized, washed and replated. For immunofluorescence, cells were plated on poly-D-lysine coated coverslips 48 h prior to fixation in 4% paraformaldehyde for 20 min. For fluorescence microscopy, slides were permeabilized for 5 min at 4ºC with 0.3% Triton X-100, and DAPI stained. Coverslips were mounted and cells were observed on a Leica DMI3000B inverted microscope and Leica AF6000E software (Leica Microsystems Inc.). A 63X/1.4 oil immersion objective was used for confocal microscopy studies on a TCS SP5 microscope (Leica) using the LAS AF software. Images were taken with the DAPI filter to eliminate bias selection of aggregates. Cells positive for cytoplasmic aggregation were quantified using ImageJ.

### Antibodies

Immunoblots and indirect immunofluorescence were performed with: rabbit polyclonal anti-TDP-43 (ProteinTech 10Z82-2-AP), anti-TDP-43 phosphorylated at Ser409/410 (Cosmo Bio USA, CAC-TIP-PTD-M01), HRP-conjugated goat anti-rabbit (Fisher Scientific PI-31460).

## Supporting information

## ACKNOWLEDGEMENTS

We thank the CNADC at Northwestern University and Eileen Bigio for providing the patient tissue; Heather True-Krob and lab members for help in setting up SDD-AGE assays; and Alessandro Vindigni for careful and critical reading of our manuscript. This work was supported by PRF, Saint Louis University (230230), ALS Association and ALS Finding a Cure and by National Institutes of Health, National Institute of Neurological Disorders and Stroke (K01 NS082391). This study was supported in part by an Alzheimer’s Disease Core Center grant (P30AG013854) from the National Institute on Aging to Northwestern University, Chicago Illinois.

