## Supplementary material for "TDP-43 oligomers detected as initial intermediate species during aggregate formation"

1. Supplementary Table
2. Supplementary Figure Legends

**Supporting Table** List and sequence of oligonucleotides used for cloning and mutagenesis.

| Name | Sequence |
|---|---|
| Bam_102_FW | 5'-CGCGGATCCAAAACATCCGATTTAATAG-3' |
| Sac_269_RV | 5'-GATCGAGCTCCTACTGTCTATTGCTATTGTG-3' |
| A315T_FW | 5'-GATGAACTTTGGTACGTTCAGCATTAATC-3' |
| A315T_RV | 5'-GATTAATGCTGAACGTACCAAAGTTCATC-3' |
| M337V_FW | 5'-CAGAGCAGTTGGGGTATGGTGGGCATGTTAGCCAGCCAG-3' |
| M337V_RV | 5'-CTGGCTGGCTAACATGCCCACCATACCCCAACTGCTCTG-3' |
| Hind_HA_mcherry_F | 5'-GACTGAAGCTTGCAATGTACCCATACGATGTTCCCGACTAC GCCGTGAGCAAGGGCGAGGAGGATAAC-3' |
| BamH_mCherry_R | 5'-CATGCGGATCCCTTGTACAGCTCGTCCATG-3' |
| BamH_TDP_FW | 5'-CATGCGGATCCTCTGAATATATTCGGGTAACCG-3' |
| Not_TDP_RV | 5'-AAGGAAAAAAGCGGCCGCCTACATTCCCCAGCCAGAAGAC-3' |
| NLS1_FW | 5'-AACTATCCAAAAGATAACGCCGCTGCGATGGATGAGACAG ATGCTTC-3' |
| NLS1_RV | 5'-GAAGCATCTGTCTCATCCATCGCAGCGGCGTTATCTTTTGG ATAGTTG-3' |

### Supplementary Figure Legends

**Figure S1. Production of soluble recombinant TDP-43.** (A) Recombinant TDP-43 expressed in bacteria purified on a nickel-charged affinity column. Representative SDS-PAGE and Coomassie blue staining of the purification samples. Arrow points to the eluted rTDP-43 fraction. (B) Removal of the SUMO tag following expression carried out on SUMO-TDP-43 immobilized on nickel-charged beads. Recombinant His-tagged Ulp1 protease was added to the beads and rTDP-43 (arrow) was recovered from the supernatant as shown in the representative Coomassie blue stained SDS-PAGE. (C) Fluorescence-based assay to measure TDP-43 RNA binding affinity for GU-repeats and estimate the active concentration of TDP-43. Representative plot of the intrinsic fluorescence (F) of recombinant TDP-43 upon titration of $(UG)_6$ RNA normalized by the fluorescence in the absence of RNA ($F_0$). Shown are F/ $F_0$ values at four different TDP-43 concentrations and data were analyzed as previously described [21]. The best fit values for the apparent equilibrium dissociation constant, $K_{d,\mathrm{app}}$ is 2.6 ± 0.5.

**Figure S2. Effect of reducing agents on the formation TDP-43 oligomers and high molecular weight aggregates detected by SDD-AGE.** Aggregation assays of TDP-43 analyzed by SDD-AGE/immunoblotting. (A) The crowding agent 0.7% PEG was added to the aggregation reaction to determine its effect on complex formation and rate of assembly. Markers to the left show the approximate molecular weight in kDa obtained from the migration of a pre-stained molecular weight marker. (B) Aggregation assays were carried out in the absence of the SUMO tag, as in Fig. 1A. (C) Aggregation assays carried out in the presence TCEP throughout purification and supplemented at day 5 (reducing). Arrows point to the time of addition of extra TCEP. The assays were also conducted under non-reducing conditions using protein purified in the presence of the short-lived reducing agent β-ME. (D) Day 3, 7 and 14 aggregation samples prepared in the presence of TCEP, as in Fig. 1A, were treated with high concentrations of reducing agents (250 mM β-ME, DTT, and TCEP). All blots are representative of > 3 individual experiments.

**Figure S3. Cellular reporter for TDP-43 aggregation HEK-TDP-43$^{NLS}$.** (A) mCherry-tagged TDP-43 transgene stably expressed in HEK293 carrying amino acid substitutions to disrupt the nuclear localization sequence (NLS). Basic residues (Red) were changed to Ala. (B) Induced expression of mCherry-TDP-43$^{NLS}$ visualized by fluorescence microscopy using DAPI to highlight nuclei. (C) Cells treated with 10 mM MG-132 for 16 hours and 0.5 mM sodium arsenite (NaAsn) for 30 minutes result in cytoplasmic aggregates which are recognized by the TDP-43 pathology marker pSer409/410. Overlay shows additional merging with DAPI. (D) Higher magnification of stress treated cells containing cytoplasmic aggregates. Scale bar, 10μm. (E) Lysate fractionation from control and MG-132/NaAsn treated cells into RIPA soluble and insoluble pellet resuspended in urea. Urea sample is 5-fold more concentrated than Total and RIPA samples. Samples loaded on SDS-PAGE and immunoblot probed with TDP-43 antibody.

**S. Figure 4. TDP-43 seeding in HEK-TDP-43$^{NLS}$ causes formation of pathologically relevant aggregates.** Immunofluorescence of HEK-TDP-43$^{NLS}$ cells transfected with rTDP-43-derived aggregates result in mCherry-positive aggregates. These inclusions are recognized by the TDP-43 pathology marker pSer409/410. Overlay shows additional merging with DAPI. (B) Higher magnification of stress treated cells containing cytoplasmic aggregates. Scale bar, 10 μm.

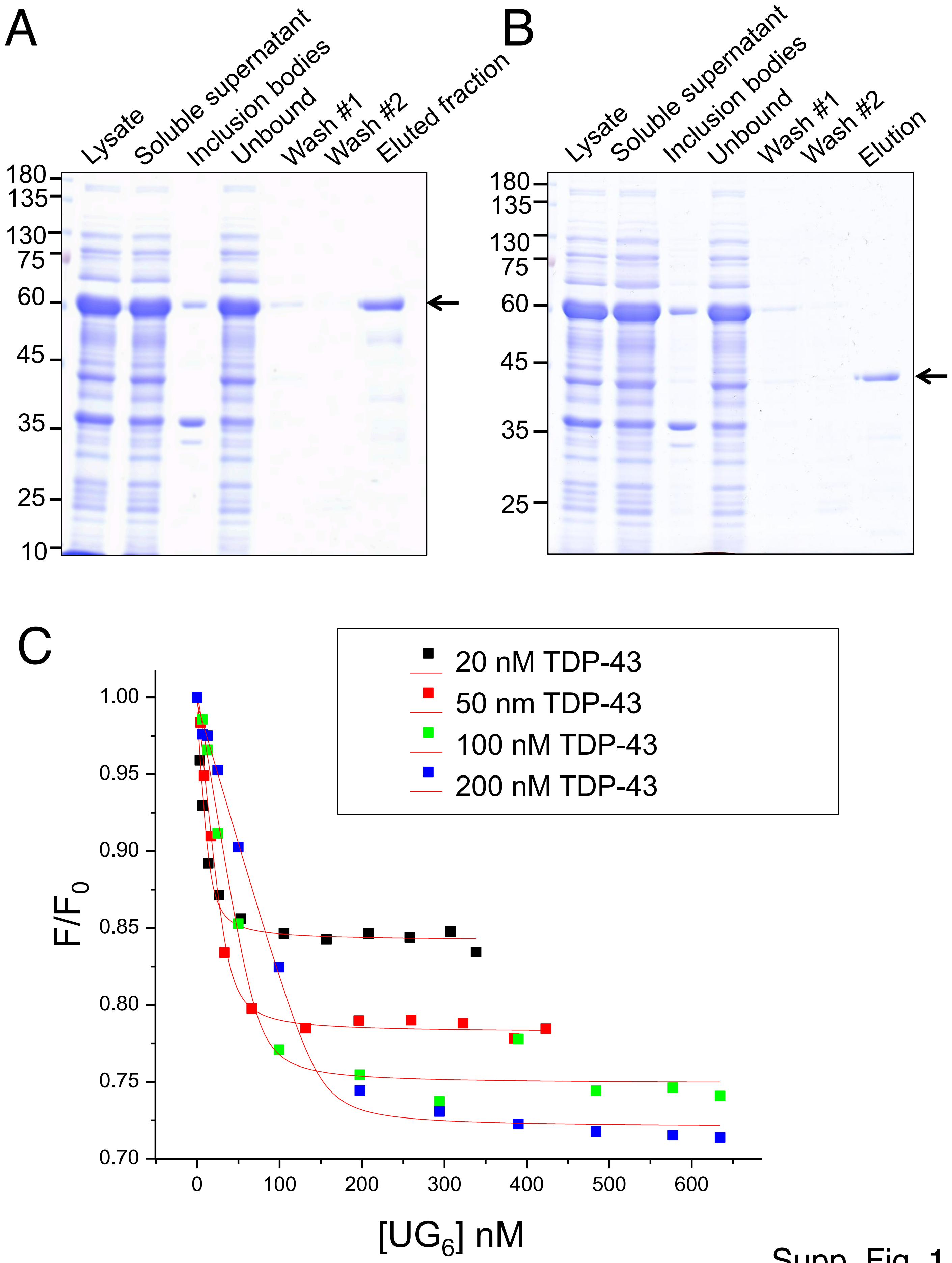
A
Lysate
Soluble supernatant
Inclusion bodies
Unbound
Wash #1
Wash #2
Eluted fraction
180
135
130
75
60
45
35
25
10
B
Lysate
Soluble supernatant
Inclusion bodies
Unbound
Wash #1
Wash #2
Elution
180
135
130
75
60
45
35
25
C
20 nM TDP-43
50 nm TDP-43
100 nM TDP-43
200 nM TDP-43
1.00
0.95
0.90
0.85
0.80
0.75
0.70
$F/F_0$
0
100
200
300
400
500
600
$[UG_6]$ nM

Supp. Fig. 1

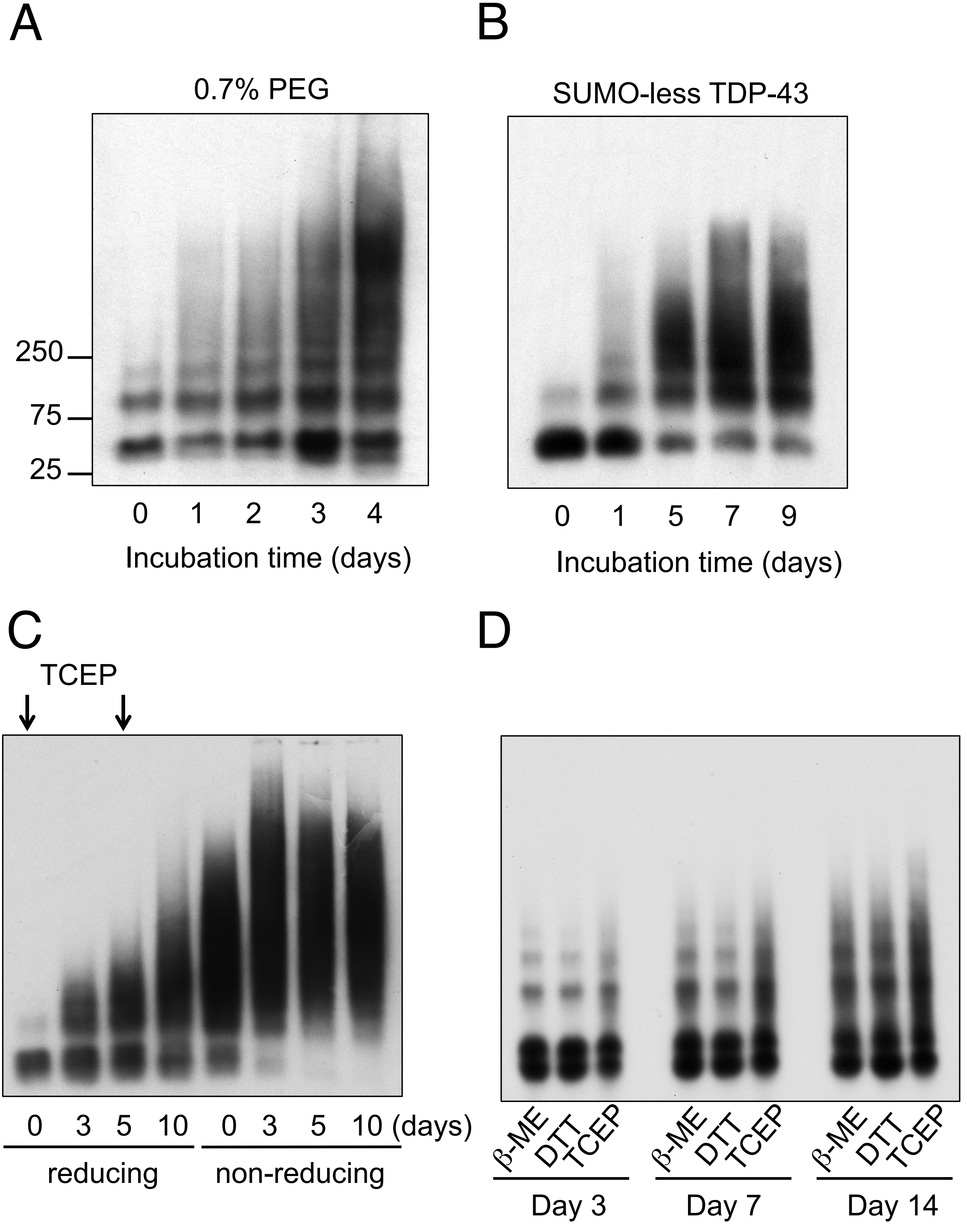
A
0.7% PEG
250
75
25
0 1 2 3 4
Incubation time (days)
B
SUMO-less TDP-43
0 1 5 7 9
Incubation time (days)
C
TCEP
0 3 5 10 0 3 5 10 (days)
reducing
non-reducing
D
β-ME DTT TCEP
Day 3
β-ME DTT TCEP
Day 7
β-ME DTT TCEP
Day 14

Supp. Fig. 2

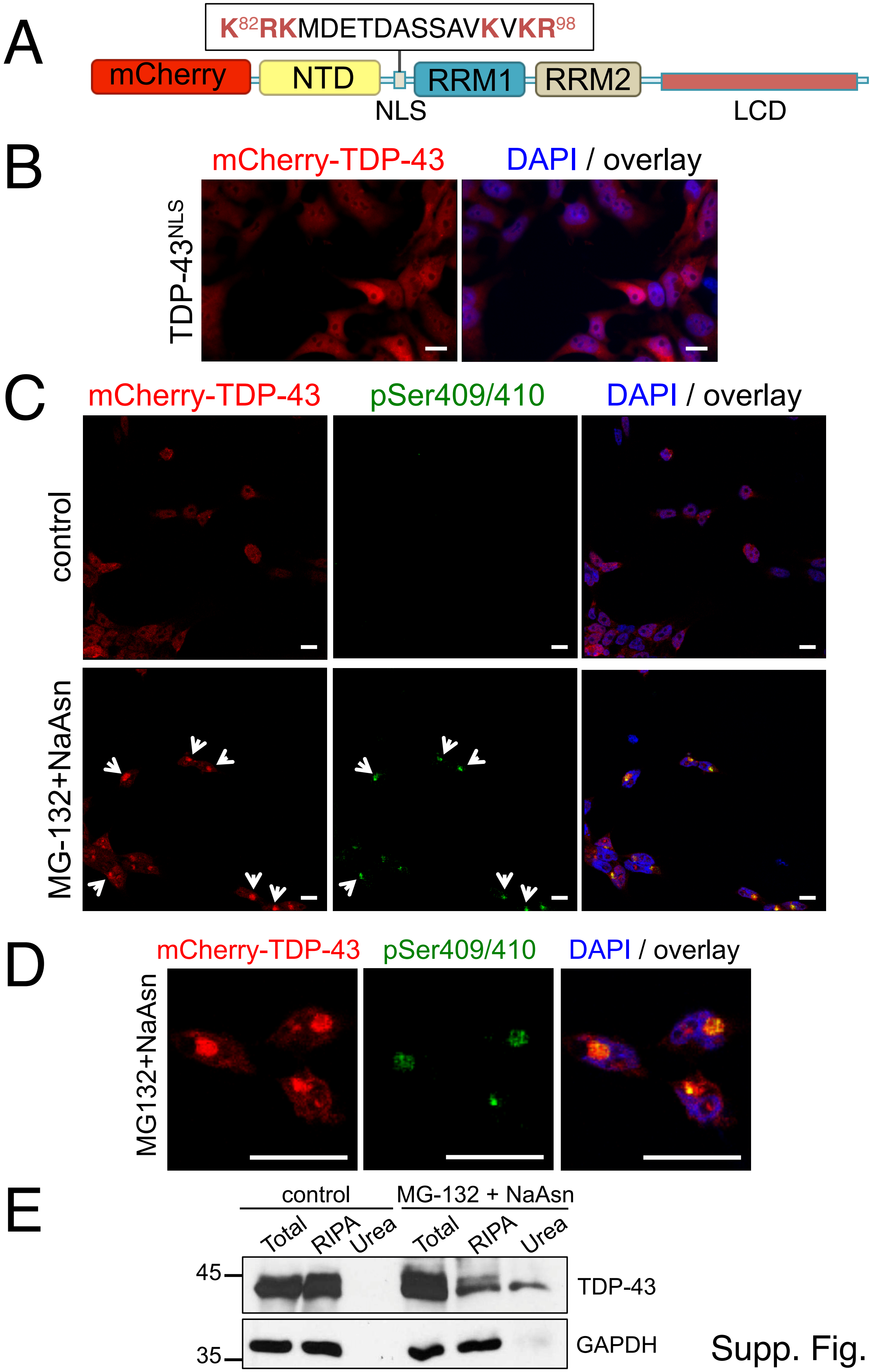
A
K82RKMDETDASSAVKVKR98
mCherry
NTD
NLS
RRM1
RRM2
LCD
B
mCherry-TDP-43
DAPI / overlay
TDP-43NLS
C
mCherry-TDP-43
pSer409/410
DAPI / overlay
control
MG-132+NaAsn
D
mCherry-TDP-43
pSer409/410
DAPI / overlay
MG132+NaAsn
E
control
MG-132 + NaAsn
Total
RIPA
Urea
Total
RIPA
Urea
45
35
TDP-43
GAPDH

Supp. Fig. 3

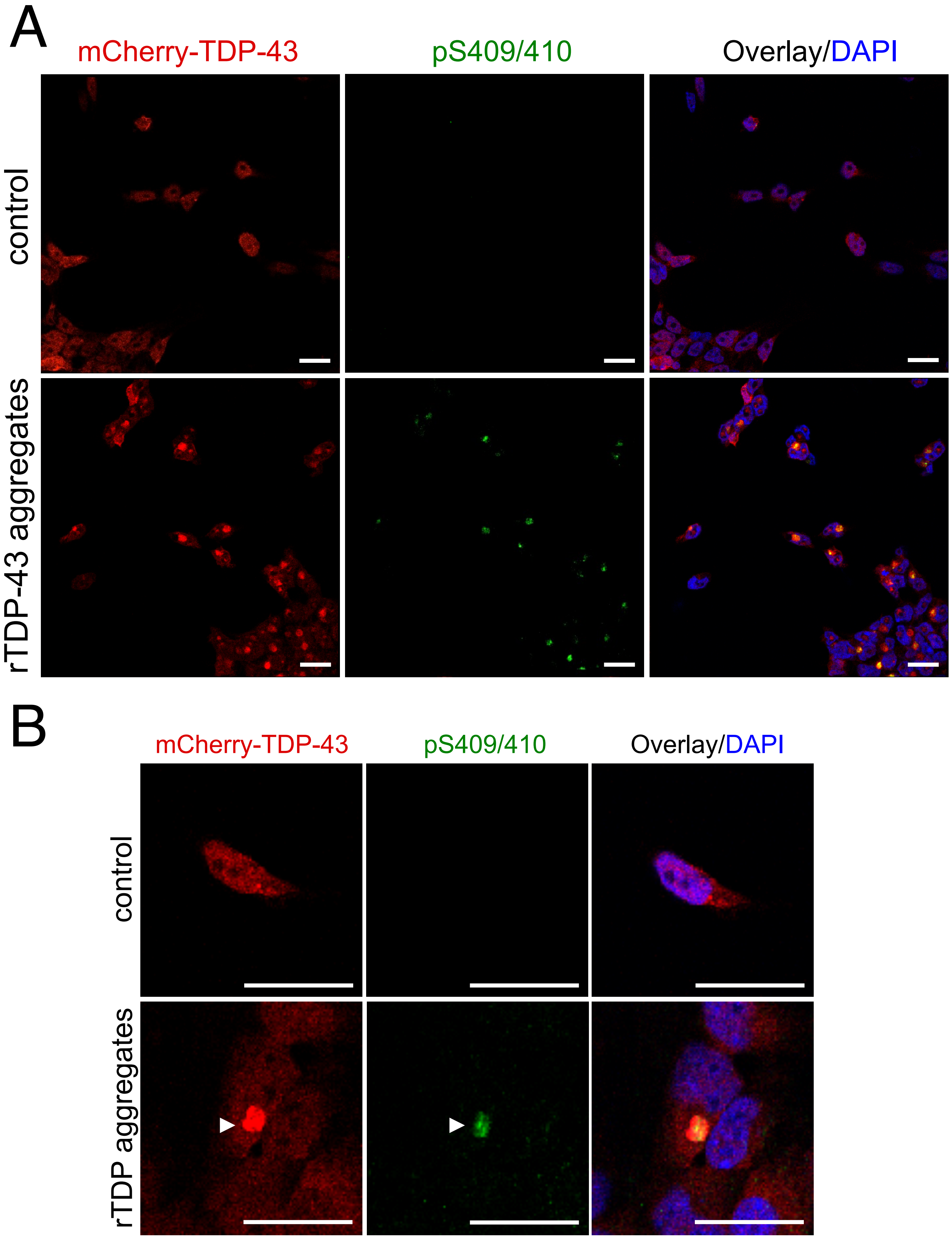
A
mCherry-TDP-43
pS409/410
Overlay/DAPI
control
rTDP-43 aggregates
B
mCherry-TDP-43
pS409/410
Overlay/DAPI
control
rTDP aggregates

Supp. Fig. 4
